# Mitogenomic Diversity in Sacred Ibis Mummies sheds light on early Egyptian practices

**DOI:** 10.1101/610584

**Authors:** Sally Wasef, Sankar Subramanian, Richard O’Rorke, Leon Huynen, Samia El-Marghani, Caitlin Curtis, Alex Popinga, Barbara Holland, Salima Ikram, Craig Millar, Eske Willerslev, David Lambert

## Abstract

The ancient catacombs of Egypt harbor millions of well-preserved mummified Sacred Ibis (*Threskiornis aethiopicus*) dating from ∼600BC. Although it is known that a very large number of these ‘votive’ mummies were sacrificed to the Egyptian God Thoth, how the ancient Egyptians obtained millions of these birds for mummification remains unresolved. Ancient Egyptian textual evidences suggest they may have been raised in dedicated large-scale farms. To investigate the most likely method used by the priests to secure birds for mummification, we report the first study of complete mitochondrial genomes of 14 Sacred Ibis mummies interred ∼2500 years ago. We analysed and compared the mitogenomic diversity among Sacred Ibis mummies to that found in modern Sacred Ibis populations from throughout Africa. The ancient birds show a high level of genetic variation comparable to that identified in modern African populations, contrary to the suggestion in ancient hieroglyphics (or ancient writings) of centralized industrial scale farming of sacrificial birds. This suggests a sustained short-term taming of the wild Sacred Ibis for the ritual yearly demand.

## Introduction

Mummification is a hallmark of ancient Egyptian civilisation and was practised on many animal species besides humans [1]. Mummies provide a unique view into the past and are potentially valuable sources of ancient DNA (aDNA). However, unfavourable environmental conditions, such as high temperatures, elevated humidity and extreme alkalinity [1-3], have resulted in debates over the authenticity of genetic results from ancient Egyptian human remains[1] [4-6]. Studies of non-human mummies have significant advantages over human mummies, since contamination is much easier to detect and control in the former. Furthermore, non-human mummified remains, particularly birds, are more numerous than human remains and can reveal information about ancient Egyptians’ religious life and their relationship to the animal world.

Animal mummies were extremely important to the people of ancient Egypt [7]. The extraordinary number of different animal species that were mummified is evidence of this [7]. By far the most numerous bird mummies found in catacombs are those of the Sacred Ibis (*T. aethiopicus*) of which no modern populations survive in Egypt. These birds disappeared from the Egyptian lands in ∼1850 [8], centuries after the cessation of the mummification practice. Approximately ten thousand Sacred Ibis mummies were deposited annually in the Sacred Animal Necropolis at Saqqara to give a final number of ∼1.75 million birds deposited at this location [9]. Similarly, Tuna el-Gebel contains approximately four million Sacred Ibis mummies, the largest known number of birds [10].

Two types of Sacred Ibis mummies have been identified [7]. One type were birds sacrificed in their millions to Thoth, the Egyptian god of wisdom and writing (Figure 1A), as ‘votive’ offerings to fulfil a prayer (Figure 1B and 1D) [7],[11]. The other type originated from ibis living in temples and were worshipped as divine incarnations of Thoth. These were mummified after their natural death [7]. There are very few sacred mummies compared to the votive ones. The latter are stacked, floor to ceiling along kilometres of catacombs at major historical sites in Egypt (Figure 1c, Figure 2a) [7]. Offering votive Sacred Ibis mummies was believed to be common practice between the Twenty-Sixth Dynasty (664-525 BC) and early Roman Period (AD 250) [12]. Radiocarbon dating [13] has established that this practice peaked between 450 and 250 BC, a result confirmed by other studies [14].

**Fig. 1.**
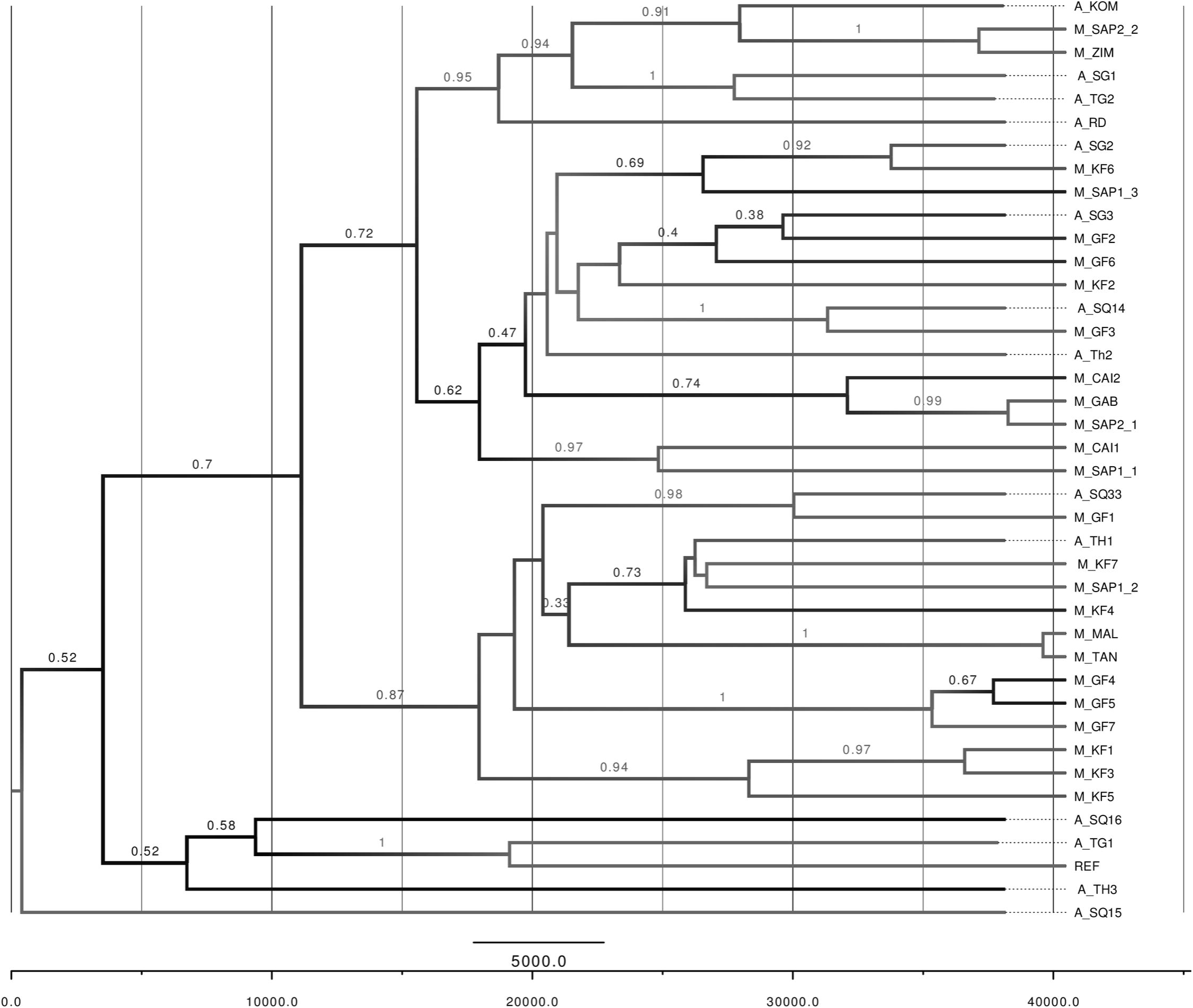
Mummified Sacred Ibis. **A** Scene from the Books of the Dead (The Egyptian museum) showing the ibis-headed god Thoth recording the result of the final judgement. **B** and **D** Example of the millions of votive mummies presented as offerings by pilgrims to the God Thoth. **C** Pottery jars containing ‘votive’ mummies stacked in the North Ibis catacomb at Saqqara.

**Fig. 2.**
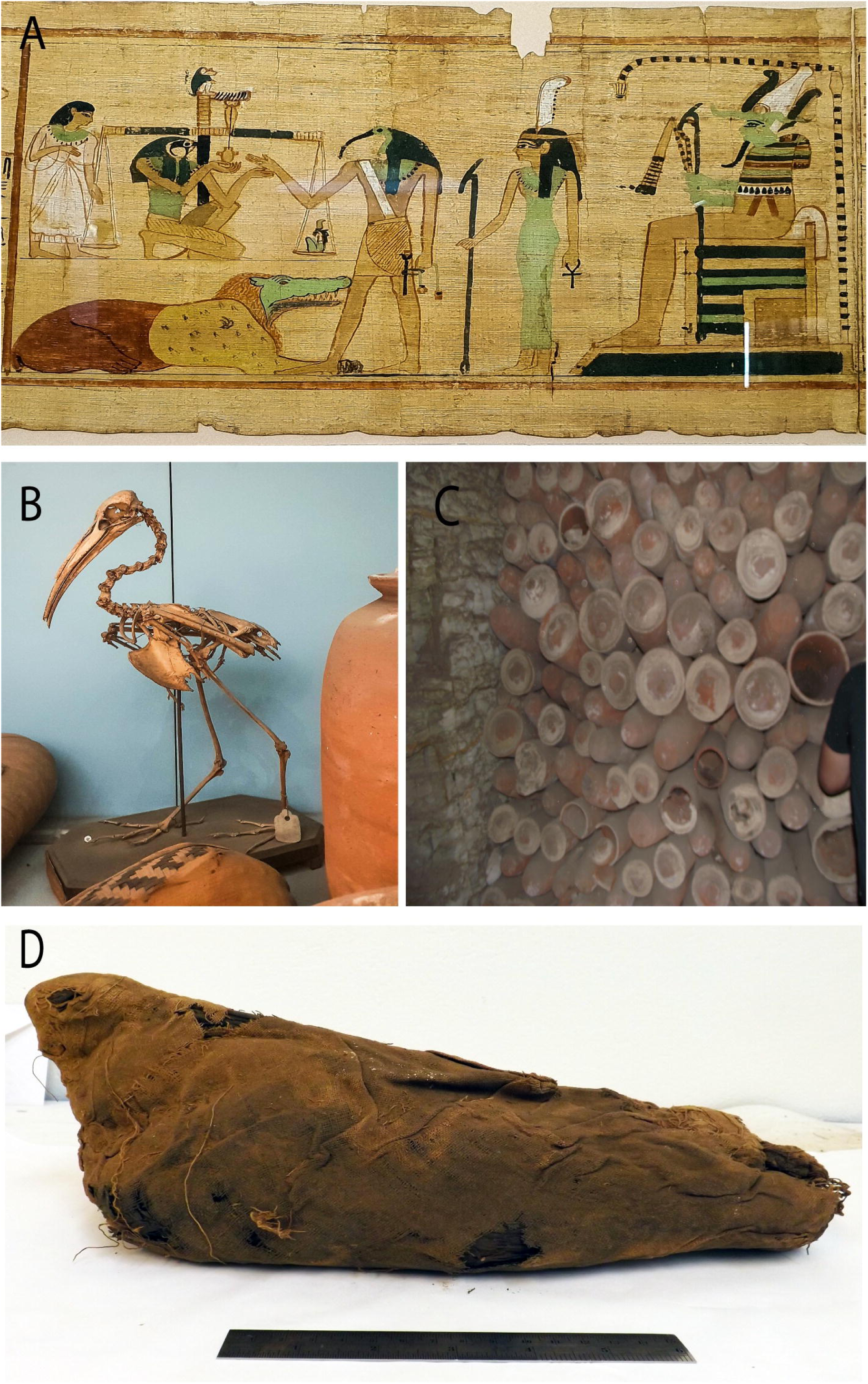
**A** Location of ancient catacombs sampled. Modern populations sampled, brown shading indicates the current distribution of Sacred Ibis. **B** Median-joining network derived from modern (orange shades) and ancient (purple shades) mitochondrial genome sequences. Circle size indicates number of samples. REF represents the Sacred Ibis mitochondrial reference genome shown in pink. Samples taken from captive Ibis at the Cairo Zoo are shown in red. **C** Principal Coordinates Analysis of distances between aligned mitogenomes of ancient (triangles) and modern (circles) Ibis. The ordination captures a very high proportion of variance in genetic distances (78.4%), with axis 1(horizontal) representing 63.2% and 15.2% for axis 2 (vertical). The asterisk denotes the reference sequence and the crosses denote Cairo Zoo. Colours in B and C correspond to the locations in A.

There is a paucity of information about how Egyptians obtained such extraordinary numbers of Sacred Ibis for sacrifice and mummification. Archaeological and ancient textual evidence [15] indicates that ancient Egyptians reared ibis on industrial scales in long-term dedicated facilities [7] [11] next to, or within temple enclosures [16]. This could be interpreted as domestication or controlled breeding. This suggestion is supported by the writings of the priest and scribe Hor of Sebennytos, from the second century BC [9]. He wrote of regularly feeding ∼60,000 Sacred Ibis with “clover and bread” [9]. It has been suggested that from the Late Period onward centralised farms provided pilgrims with Sacred Ibises that could be mummified and offered at Thoth temples [17, 18].

The early presence of Sacred Ibis mummies at Tuna al-Gebel were thought to have been sourced from all over Egypt as indicated by the demotic writings (ancient Egyptian type of writings) (Figure 3) which were found accompanying the mummy wrappings, papyri, or jars [19]. Texts recording the donor, date, and provenance of birds indicate that Sacred Ibis mummies, sometimes accompanied with eggs, or even separate bundles of eggs, were sent to Tuna el-Gebel from other locations. (Figure 3) [17-19]. It appears that it was not only main cities like Aswan, Ptolemais-Psois, Hermopolis or Heliopolis that provided Sacred Ibis to Tuna, but also smaller sites which have not yet been located [19]. Important information on how the mummified Sacred Ibis were transferred from El-Fayoum region to Tuna al-Gebel has also been recorded on papyri [17] and it is believed that these transfers continued into late Ptolemaic times. The papyri found inside the jars in some of the subterranean galleries date to the time of the Persian ruler Darius I (522-486 B.C) and record the transportation of mummified Sacred Ibises and their subsequent offering at Tuna al-Gebel [17] in the south. Based on textual evidence found with buried mummies (Figure 3), sending mummified Sacred Ibises from numerous other Egyptian locations to Tuna al-Gebel continued after 305 BC.

**Fig. 3.**
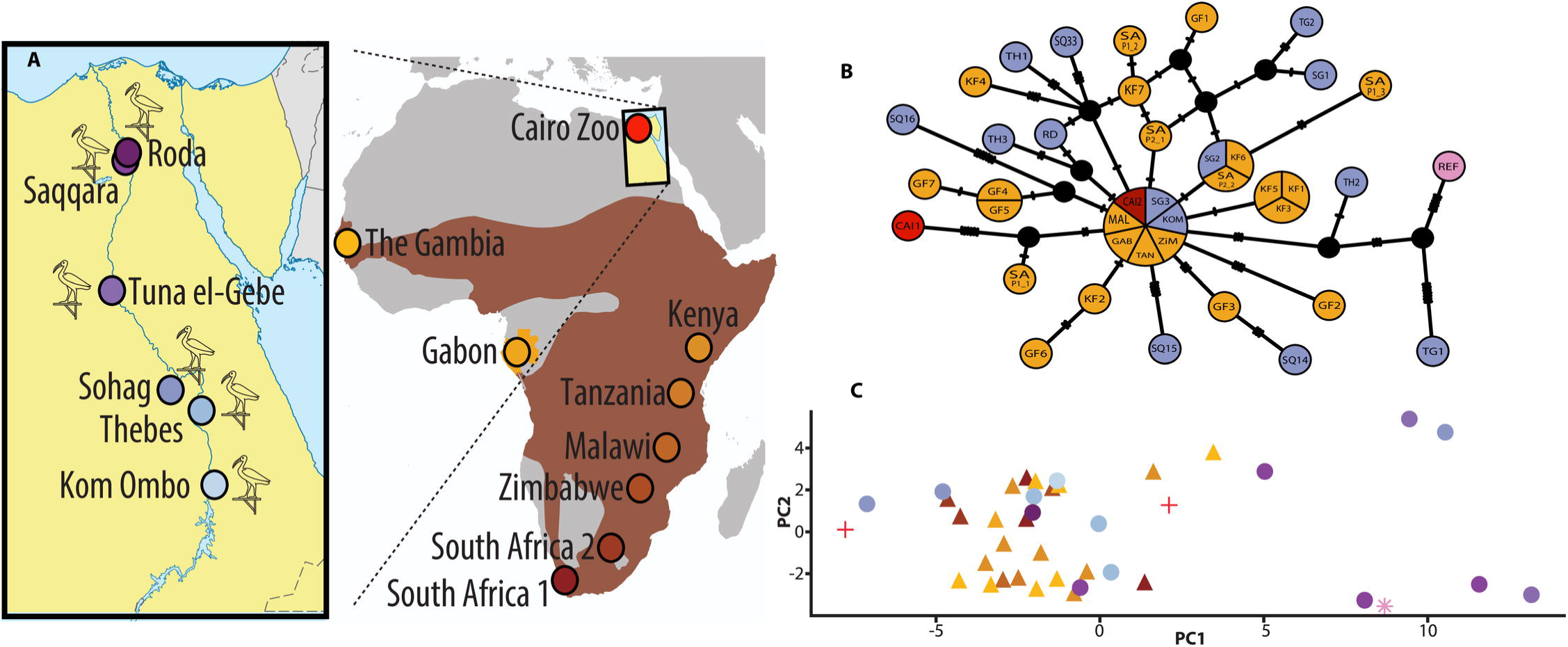
Ancient inscriptions on a pottery jar of a mummified Ibis from Tuna el-Gebel. This inscription recorded the date the mummy was offered to Thoth, by whom, where it was bought from and the name of the priest. From Mahmoud Ebeid, BIFAO 106 (2006), p. 57-74.

By the Ptolemaic period, the demand for Sacred Ibis mummies intensified, leading to a more localised system, rather than depending on transfers from all over Egypt to the main burial necropolis [18]. By this time the nationwide transfer of birds became limited to the sacred or the ‘ritual’ type of the Sacred Ibis, those were kept in the temple as representation of Thoth. During the reign of King Ptolemy I (c. 367BC – c. 283 BC), villagers were forced to both work and pay for the support of Sacred Ibis farming (ibiotropheia), which led to the presence of approximately a dozen Sacred Ibis breeding farms in the area of Hermopolis. Although it is unknown if the birds were sourced every year from the wild and tamed, or if they were bred in captivity over generations, these farms were equipped to raise birds and were surrounded with fields that supplied Sacred Ibis colonies with cereals [18].

Evidence that Egyptian mummified ibises were raised in captivity stems as far back as 1825, from the French naturalist Georges Cuvier. Describing an ibis mummy from Thebes that he had unwrapped to study, Cuvier noted: “One sees that this mummy must have come from a bird held in domesticity in the temples, for its left humerus was broken and reset. It is probable that a wild bird which had had its wing broken would have perished before being healed, for lack of being able to chase its prey or to escape its enemies” [20].

During the Ptolemaic era, the level of production of each of the local Hermopolitan Sacred Ibis’ farms has been estimated to be around a thousand mummies annually. Kessler [21] proposed the existence of around fifteen local ibiotropheia producing an estimate of fifteen thousand mummies, which were brought to Tuna al-Gebel each year [18].

Sacred Ibis eggs were collected during the Saite period (664 BC – 525 BC) from breeding places and wild colonies and were sent to Tuna al-Gebel together with wrapped mummies [18]. Some scholars hypothesized that these might have come from an artificial breeding hatchery, although no hard evidence has been found to support this suggestion [22].

Alternatively seasonal taming of wild birds has been suggested [7] where votive mummies were reared (but not domesticated) by priests in natural habitats close to the temples [9, 18]. This is thought to have occurred in locations such as ‘the Lake of the Pharaoh’, known later as the Lake of Abusir located between Abusir and Saqqara [9], and ‘the swamp’ near Tuna el-Gebel. The swamp probably refers to a natural basin that filled annually with the Nile inundation [19]. Furthermore, in the Ptolemaic period it has been reported that mummies were rarely sent from across Egypt to Tuna el-Gebel, but instead, ten to fifteen local Sacred Ibis breeding sites near Tuna el-Gebel’s appeared to supply this temple [18].

## Materials and Methods

### Materials

With the permission of the Egyptian Ministry of State for Antiquity, samples were collected from the three main Ibis catacombs: Saqqara, Tuna El-Gebel, and Sohag (Abydos), where they were retrieved from the storage magazine at Sohag (Figure S1). Also, a number of museums worldwide (Table 1) agreed to send ancient Ibis tissues for this research. Blood and feather samples from contemporary Sacred Ibis were collected from various locations across Africa (Table 1). No ethics clearance was requested for the collection of the Sacred Ibis feathers as no harm was involved in the process. In the case of blood samples from South Africa these were collected under an Animal Ethics approval from the University of Cape Town as detailed below in the ethics section.

**Table 1:**
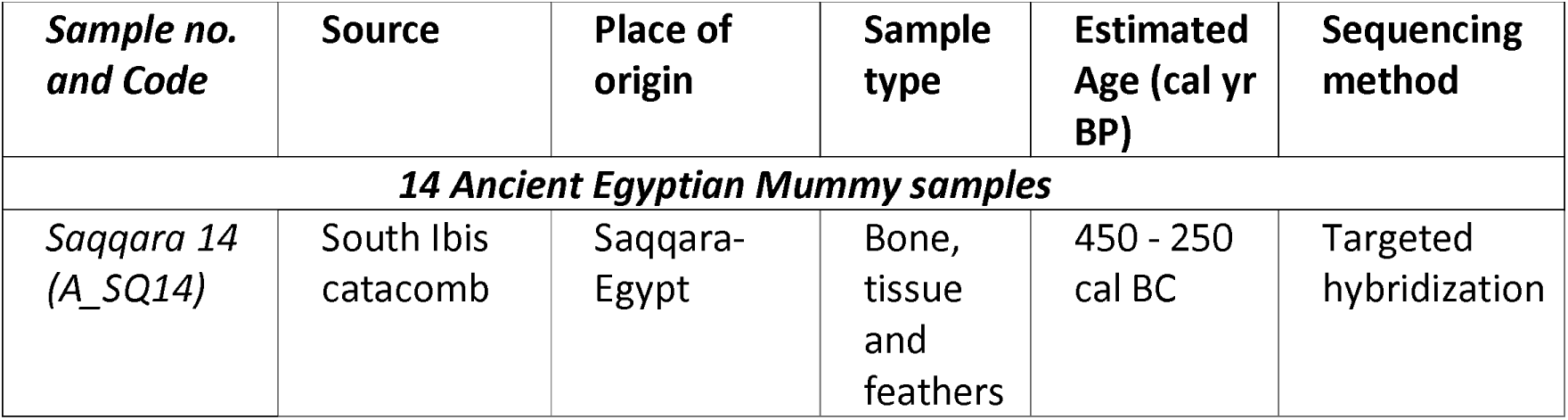
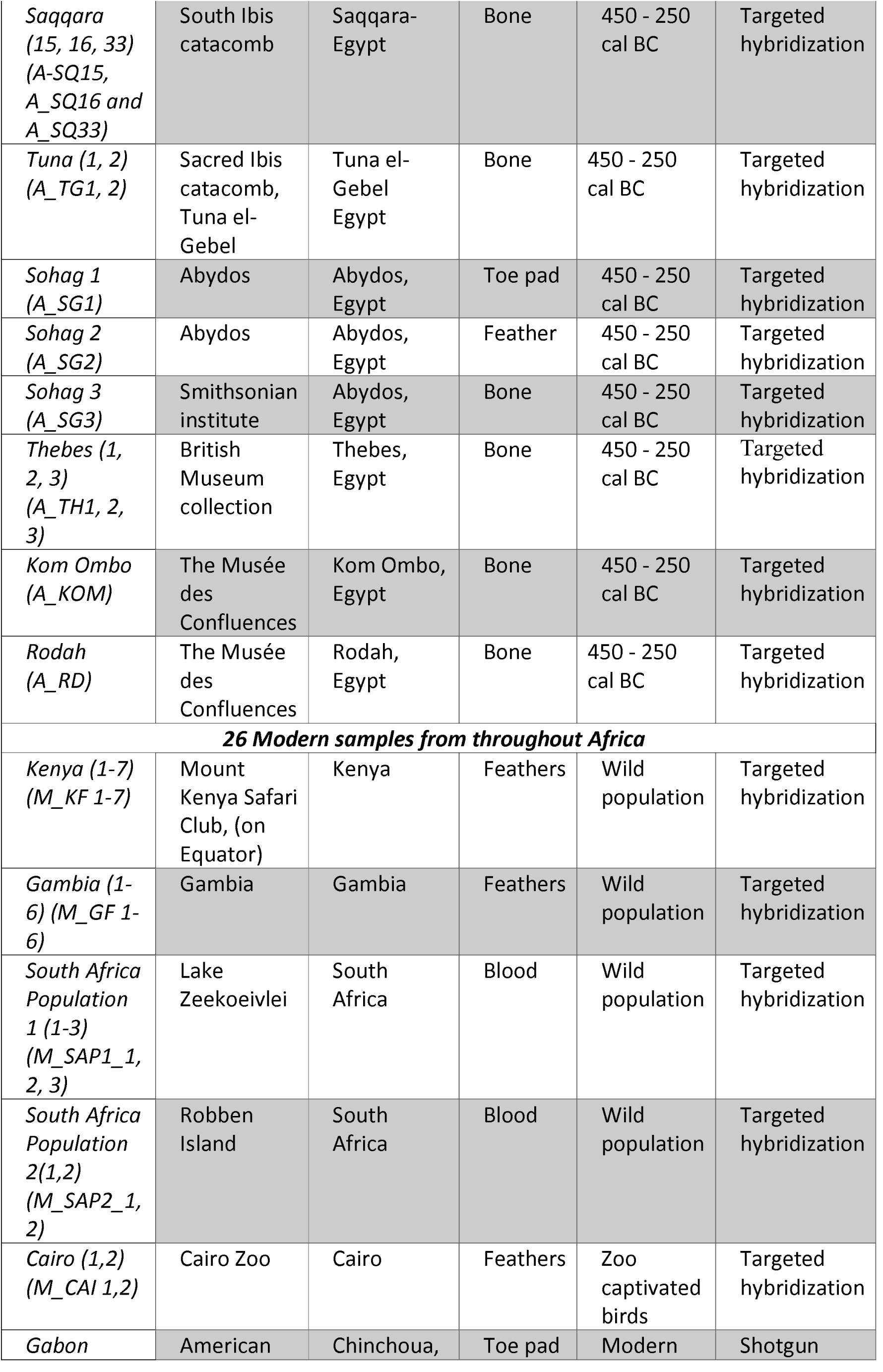
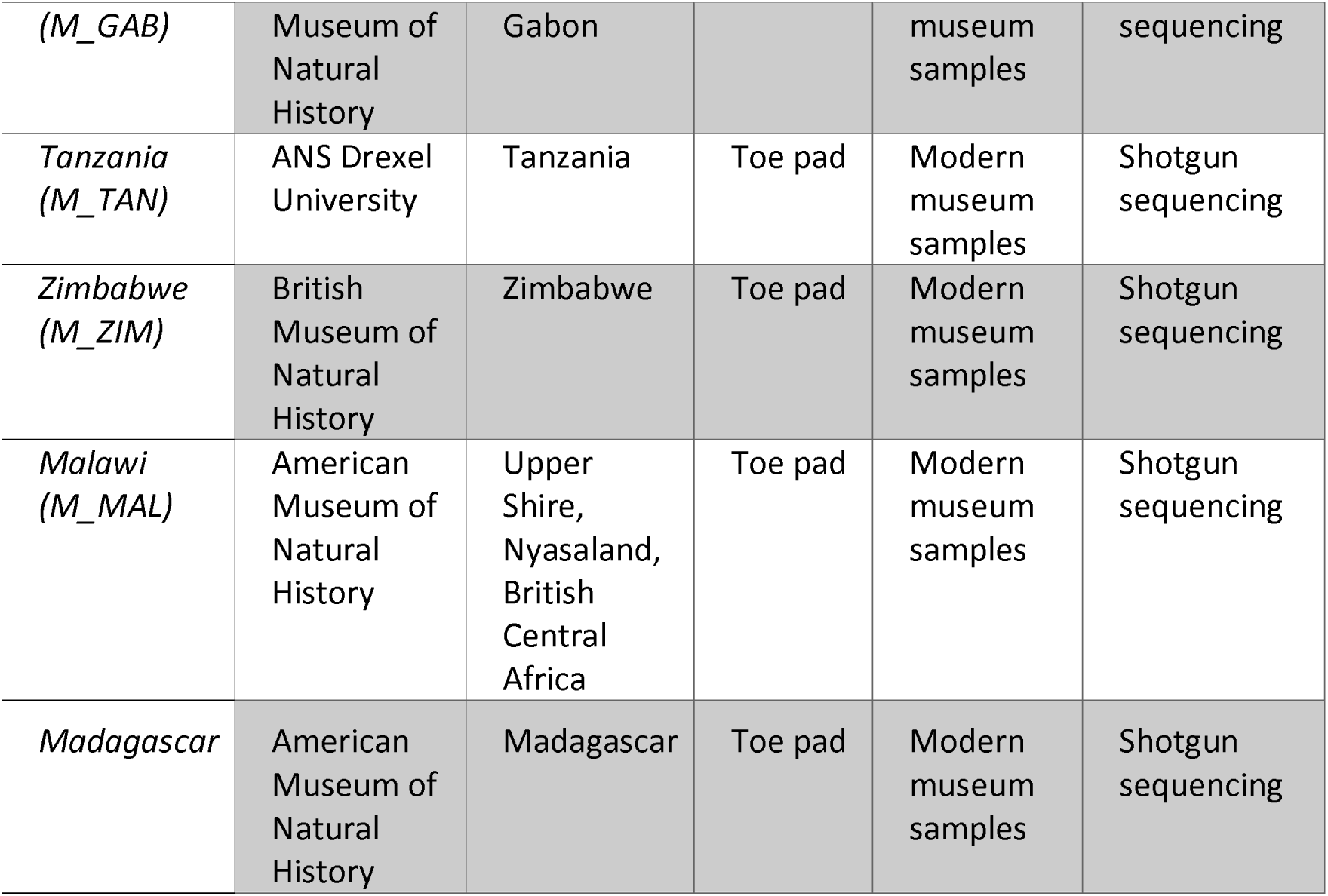
Sacred Ibis samples. Details of the location, tissue type and estimated ages of both modern and ancient Sacred Ibis samples sequenced in this research project. Age reported in the table is an estimated age based on the samples reported in Wasef et al., 2015 [13] from same locations.

## Methods

### Modern DNA Extraction

DNA was extracted from modern Sacred Ibis blood and feather samples by incubation, with rotation, overnight at 56°C, with 5 uL of blood or part of the feather including its quill, in SET buffer containing 10% SDS, 20 mg/mL proteinase K, and 50mM DTT. The mix was then extracted with an equal volume of phenol:chloroform:isoamyl alcohol (25:24:1), followed by a chloroform step and finally purified using a Qiagen MinElute column as outlined by the manufacturer.

### Ancient DNA Extraction

Sacred Ibis mummies were commonly preserved by being dipped in melted resin or salted and resined [23]. This can be detrimental to the recovery of DNA because this process initiates an oxidation reaction that burns the bones from the inside, leaving only powder inside the wrapping [18]. Luckily, not all the Sacred Ibis were mummified in this way and some mummies were found in a well-preserved state with feathers and tissue still largely intact. DNA extractions and further processing were carried out in dedicated ancient DNA facilities at the Al Kasr Al Ani Medical School in Cairo and at Griffith University, Nathan, Australia. Prior to extraction, ancient bone, feather or tissue samples were cleaned with 10% bleach and then 80% alcohol. The outer bone layer was then removed and the remaining bone fragments were crushed to a fine powder. Approximately 50 mg of bone powder was used for extraction according to the method of Dabney et al., 2013 [24], with minor changes such as the addition of 20 uL of 50mM DTT and incubated overnight at 40°C with rotation. Next a 10×PB buffer (Qiagen) was added to the extract [24] and the DNA was purified using Qiagen DNeasy Blood & Tissue Kit columns, as recommended by the manufacturer. DNA was eluted from the column using 50 μL of Ultrapure water. For ancient toe pads, tissue, and feathers, the samples were sliced with a scalpel, and extracted with 200 uL of SET buffer, 40 uL of 10% SDS, 20 uL of 20 mg/ml Proteinase K and 20 uL of 50mM DTT and the rest of steps to clean DNA are as outlined for ancient bone samples.

### Construction of Illumina sequencing libraries

Illumina sequencing libraries were constructed from modern DNA using a NEBNext Ultra™ DNA Library Prep Kit (NEB) as described by the manufacturer. The resulting library was amplified by PCR for 15 – 22 cycles using Phusion® High-Fidelity PCR Master Mix in GC Buffer (NEB) and a NEbNext Universal PCR Primer for Illumina and an Illumina multiplex index primer. Amplified libraries were purified using AMPure XP beads for library clean-up. Ancient DNA Illumina Sequencing libraries were constructed using a modification of the method of Meyer and Kircher [25]. KAPA HiFi Uracil+ polymerase Master Mix (KAPABiosystems) was used for the ancient DNA libraries PCR amplification for between 14 to 20 cycles. Amplified libraries were cleaned up with 1x AMPure XP beads.

### Quality control

Modern DNA extractions from blood were visualized on the Agilent Bioanalyzer 2100 as an approximate check for DNA size before and after using the Covaris for shearing the DNA and the AMPure XP beads for size selection. Ancient DNA extracts that showed bacterial or modern contamination in the form of high molecular weight product were excluded. Amplified libraries were visualised using High-Sensitivity DNA chips on an Agilent Bioanalyzer 2100; to adjust for the optimal number of PCR cycles with the minimal percentage of PCR clonal sequences; for the insert size of the library; and to determine the required amount of each library for either direct sequencing or target capture hybridization.

### Target capture hybridisation

Capture baits to the complete Sacred Ibis mitochondrial genome were designed as single stranded 80-mer biotinylated RNAs with 5 base overlaps by MYcroarray [26, 27]. The capture enrichment was performed for both amplified Illumina libraries for both ancient and modern samples, according to the manufacturer protocol using hybridisation temperature of 55°C-65°C for 2-3 days [26, 27] and amplified for 10-18 cycles using Phusion® High-Fidelity PCR Master Mix and GC Buffer (NEB).

### Illumina second-generation sequencing

Purified indexed libraries were sequenced either using the MiSeq sequencer at Griffith University, Brisbane, Australia or using the single-end reads for 100 cycles on the HiSeq 2500 at the Danish National High-Throughput DNA Sequencing Facility in Copenhagen.

### Bioinformatics

Sequence reads were initially processed using the fastx_toolkit V0.0.13. Adapter sequences and reads shorter than 25 bases were removed, and low-quality bases were trimmed. Following processing, reads were aligned to the Sacred Ibis mitochondrial reference genome (NC 013146.1) using BWA V0.6.2-r126 [28]. SAMtools [29] was used to extract data, index, sort, and view output files. Qualimap [30] was used to assess alignment quality. MapDamage2.0 [31] was used to estimate ancient DNA authenticity by measuring the levels of post-mortem damage [27]. The 14 ancient and 26 modern sequences were aligned using the online version of MAFFT [32]. Population genetics analyses from DNA sequence data were carried out using DnaSP v5 software [33], and was used to generate the following statistics: Haplotype diversity [34], the number of haplotypes [34] in the Sacred Ibis genomes. The entire mitochondrial genome sequences of the 40 samples in total, excluding alignment gaps were used in the network and phylogenetic construction. Median-joining networks for inferring intraspecific phylogenies were constructed with NETWORK v. 4.6.13 [35] (Figure 2b). A Bayesian estimate of the phylogeny was constructed using BEAST 2 [36] for ancient and modern Sacred Ibis complete mitochondrial genomes. This assumes a Bayesian skyline plot, a strict molecular clock and the bModelTest [36] [37] approach to averaging over site models. The x-axis is years before present, and edges are labelled with their posterior clade probabilities (Figure 4).

**Fig. 4.** A Bayesian Summary tree. The x-axis is years before present, and the edges are labelled with posterior clade probability. M_ indicates modern samples from Africa. A_ indicates Ancient Sacred ibis mummies. Samples’ code used in the tree are listed in table 1.

Mitogenomic diversity within and among populations was estimated using the Maximum composite likelihood method employed in MEGA6 [38]. We first estimated the extent of rate variation among sites (α). This was then used to estimate the diversity within and among populations. Using the exon boundary annotations for the Ibis reference mt-genome sequence, we extracted the coding sequences (CDS) for each gene and concatenated them. We used the software PAML [39] to estimate dN/dS ratios. For this purpose, we used the concatenated alignment containing 13 protein coding genes from modern and ancient mitogenomes and used only the codons present in all sequences. We estimated a single dN, dS and dN/dS for the whole tree using option ‘one’ in *codeml*. These were estimated for modern and ancient sequences separately. We used a bootstrap resampling procedure (1000 replicates) and obtained point estimates for each bootstrap replicate using PAML. These were used to estimate the standard error. A Principal Coordinates Analysis was performed on a pairwise distance matrix of aligned sequences under Kimura’s 2-parameters distance model [40], using the ape package [41] in R and visualised using the R package ggplot2 [42] (Figure 2C).

## Results

We recovered tissue and extracted DNA from 40 Sacred Ibis mummies from six Egyptian catacombs. In addition, modern mitochondrial diversity of wild ibis was determined from 26 birds sampled from 10 locations across Africa. This represents the species’ current geographic range (Figure 2A, Table 1). Twenty ancient extractions were selected for shotgun sequencing to measure levels of endogenous DNA, which was typically low (0.06%). These minute levels of endogenous DNA meant that we needed to enrich mitochondrial DNA libraries by targeted hybridisation using biotinylated RNA baits [26] designed against the Sacred Ibis mitogenome (GenBank NC_013146.1). Total sequences of the mitochondrial genome retrieved after enrichment using baits ranged from 5.3x–336x, improving mitogenome coverage to 1.5x–35x (Supplementary Table S1) [26] [27]. This resulted in the recovery of 14 complete ancient and 26 modern mitogenomes (Supplementary Table S1). We show that populations of modern African Sacred Ibis and ancient mummies show similar mitochondrial diversity patterns, as evidenced by haplotype network analyses (Figure 2B) [43] and Principal Coordinates Analysis (PCoA, Figure 2C). The levels of mitogenomic variation within ancient Sacred Ibis populations, and those within modern populations, are not significantly different (Table 2).

**Table 2:**
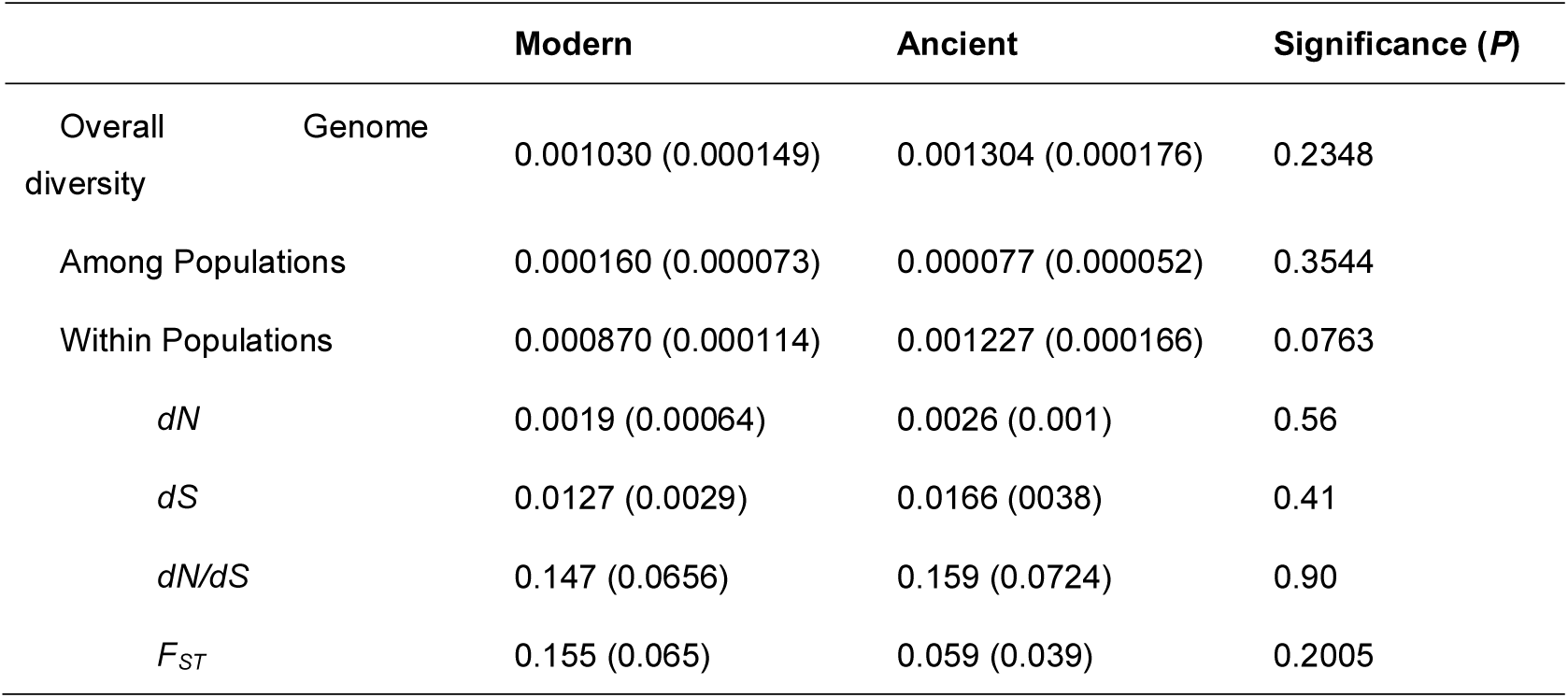
Intra and inter population pairwise comparisons of mitogenomic variation in ancient and modern Sacred Ibis. P-values > 0.05 indicate no significant difference between modern and ancient Sacred Ibis populations. Standard deviations (SD) are in brackets. A Z-test was used to obtain P-values.

Based on the Bayesian estimate of the phylogeny constructed [36], comparing a tip-dated model (used to generate the current tree in Figure 4) with an equivalent non-tip-dated model, does not reveal strong evidence for measurable evolutionary change between the ancient and the modern specimens (Figure 4). This is likely due to a lack of power and we expect that if the analysis were repeated with older ibis genomes and higher read coverage (to better detect non-damaged variants) then measurable evolution might be detected.

### Pre-capture Results

Initial shotgun sequencing of genomic libraries showed that modern feather DNA samples yielded the most endogenous mitochondrial DNA (x□= 0.04%; calculated as percentage of unique sequences versus total number of reads). This was followed by ancient toe pad samples (x = 0.01%), ancient feather and bone, modern blood samples (x□= 0.002%), and finally ancient soft tissue (x□= 0.0002%) [27]. The low amount of endogenous mitochondrial DNA detected in modern blood is likely due to the low mitochondrial DNA copy number in avian blood [44]. DNA length varied significantly amongst the ancient samples [27].

The number of duplicated sequences varied considerably amongst ancient tissues from 3.03% to 89.26% with no significant differences noted between the various tissues. Modern feathers were shown to have the least number of duplicates at 10.6 ± 9.6%, while modern blood had a high level of duplicated sequences at 82.4 ± 3.6%. Ancient samples have been shown to display clearer damage and fragmentation patterns characteristic of endogenous ancient DNA [27] [31].

### Target capture results

Enrichment rates were determined for each sample by calculating the percentage of the unique (non-clonal) sequences aligned to the mitochondrial reference genome; pre- and post-capture hybridisation enrichment (Supplementary Table S1). Our results indicated that regardless of the sample used or the hybridisation temperature, there was significant enrichment in the unique endogenous content of the captured libraries [27] (Supplementary Table S1). Also, by comparing the insert size of the ancient and modern pre-capture sequences to their equivalent post-capture sequences (Supplementary Table S1), we found a slight increase in the mean read length of the unique sequences for most samples (1.2 fold). Those results are consistent with the previous observations [45, 46]. In terms of the tested hybridisation temperature, we show that hybridisation and washing temperatures of 65°C resulted in increased enrichment of modern mitochondrial DNA (x □= 199 fold) when compared to ancient DNA. For the ancient sacred Ibis samples, our results show that the best enrichment temperature was 57°C, with enrichment rates between 54×to 4705×[27] (Supplementary Table S1).

## Discussion

The Sacred Ibis played a significant role in ancient Egypt through its representation of Thoth, the god of writing, scribes, wisdom, time and justice and as the deputy of the sun-god Horus-Re. Sacred Ibis were nurtured, bred, and mummified with the same attention to ritual detail, as was given to many humans of that time [7]. There is a large amount of archaeological evidence for Sacred Ibis in ancient Egypt, particularly in the burial grounds at Saqqara, Abydos, Tuna el-Gebel and Thebes [7]. The analysis of mitogenomic data from a number of ancient and modern Sacred Ibises allowed us to test theories proposed from archaeological studies about the farming system used by ancient Egyptian priests in order to maintain a sufficient number of Sacred Ibises to meet demand for cultic activities and has clarified the origin of one of Egypt’s iconic birds. The use of newly developed ancient DNA technologies had allowed us to further test those theories associated with ancient Egyptian civilization.

We examined the mitogenomic data of the mummified Ibises to test the centralised mass-farming production hypothesis as opposed to Sacred Ibis being sourced seasonally by short-term taming of wild individuals. If the former scenario were true, high levels of inbreeding and population bottlenecks following the mass sacrifice of birds would have led to low genetic diversity among mummies. We would also expect a higher ratio of non-synonymous to synonymous diversity (dN/dS) in ancient Sacred Ibis, compared to those from modern populations, due to the accumulation of deleterious variants. Alternatively, it is possible that birds were captured from wild populations and kept near temples in short-term seasonal farms maintained by locals under supervision of priests, or birds were regularly fed in order to attract them to the freshwater breeding sites. These alternate hypotheses imply high overall mitogenomic diversity and high inter-catacomb diversity. In contrast to the centralised-farming hypothesis, the d_N_/d_S_ ratio of ancient populations and levels of population structure would be expected to be similar to that found in modern populations. The supplementary feeding would also have enhanced and supported a large population size.

The overall mitogenome diversity observed for ancient samples was not significantly different to that observed for modern samples (Using Z test, P=0.23, Table 2). In addition, the inter-catacomb diversities among ancient populations were similar to the inter-population diversities obtained for modern populations (Table 2). Furthermore, the dN/dS ratio estimated among ancient mitogenomes of 0.159 (0.072) were not significantly different (P=0.9). The standard error (SE) was calculated using a bootstrap resampling (PAML) and the variance was used to perform a Z test to that estimated for modern populations (0.147 (0.065). The *F*_*ST*_ values estimated for ancient mitogenomes revealed very low and insignificant (P=0.20) levels of structure among the populations from different catacombs (Table 2) and provide no support for hypothetical long-term farming practices.

In summary, our rejection of long-term centralised farming is based on threefold lines of evidence: First, the overall genomic diversity of ancient Sacred Ibis populations within and among catacombs was comparable to that found in and among modern Sacred Ibis distributed throughout Africa. If breeding of ancient Sacred Ibis were conducted using a small number of founding populations, we would have expected low genetic diversity amongst the ancient Sacred Ibis compared to that of modern populations. Second, the diversity observed at evolutionarily constrained (non-synonymous) sites of protein-coding genes of ancient samples was similar to that recorded for contemporary Sacred Ibis populations. In contrast, we would expect much higher *d*_*N*_*/d*_*S*_ diversity in ancient Sacred Ibis if they were bred over extensive time frames in dedicated farms. Finally, we did not observe significant population structure among the Sacred Ibis populations from different catacombs. Together, these data suggest that the most probable scenario is that local Sacred Ibis were tended in the natural habitats or small, localised farms. If they were deliberately farmed, it is likely that this would have been for only short time periods (perhaps a single season), before being sacrificed and entombed.

## Ethics

Permission to carry out fieldwork is detailed in the methods section. Collection of modern blood samples from South Africa were carried by Doug Harebottle (Permission number: 69052850438083).

## Data accessibility

Ancient and modern Sacred Ibis DNA sequences generated in this study are available in GenBank (NC_xxxx).

## Authors Contribution

SW performed the research, analysed data, contributed new reagents / analytic tools, collected the ancient samples, wrote the manuscript, with input from all authors; SS analysed data, contributed new reagents / analytic tools, helped draft the manuscript; RO analysed data, contributed new reagents / analytic tools, helped draft the manuscript; LH performed the research, helped draft the manuscript; SEM collected the ancient samples, helped draft the manuscript; CC performed the research, acquired modern and ancient museum samples, helped draft the manuscript; AP contributed new analytic tools, helped draft the manuscript; BH analysed data, helped draft the manuscript; SI acquired modern and ancient museum samples, helped draft the manuscript; CM. designed the research, analysed data, wrote the manuscript, with input from all authors; EW designed the research, helped draft the manuscript; DL designed the research, analysed data, wrote the manuscript, with input from all authors. All authors gave final approval for publication and agree to be held accountable for the work performed therein.

## Funding

Human Frontier Science is acknowledged for financial support in the form of a grant to PIs, Lambert, Ikram, Holland, and Willerslev (RGP0036/2011). Wasef thanks Griffith University for a PhD scholarship.

## Acknowledgements

We are grateful to The Danish National High-Throughput DNA Sequencing Centre for sequencing the samples. We also thank Doug Harebottle for collecting the blood samples from South Africa; and Clive Richard Barlow for collecting the feathers from Gambia. The authors are grateful to the Ministry of Antiquities for permission to collect and conduct research on these ancient samples. We also appreciate permission from the Al Kasr Al Ani Medical School for allowing Dr Wasef to use the ancient DNA laboratory in Cairo. A number of museums kindly provided material for this study including: The British Museum, The Smithsonian institute, the Ancient Egyptian Animal Bio Bank at Manchester Museum, especially Dr. Lidija M. McKnight and the Musée des Confluences, Lyon, France, particularly Stephanie Porcier. We also thank the Academy of Natural Sciences of Drexel University (Philadelphia, PA), The American Museum of Natural History (New York, NY), and the British Museum of Natural History for providing access to modern ibis samples. We thank Professor Alexei Drummond from the University of Auckland for advice regarding phylogenetic analyses. The authors are also grateful to the Environmental Futures Research Institute and Griffith University for additional support.

## References

1. Gilbert MT, Barnes I, Collins MJ, Smith C, Eklund J, Goudsmit J, et al. Long-term survival of ancient DNA in Egypt: response to Zink and Nerlich (2003). Am J Phys Anthropol. 2005;128(1):110-4; discussion 5-8.

2. Zink AR, Nerlich AG. Long-term survival of ancient DNA in Egypt: Reply to Gilbert et al. American Journal of Physical Anthropology. 2005;128(1):115–8.

3. Zivie A, Lichtenberg R. The cats of the goddess Bastet. In: Ikram S, editor. Divine Creatures: Animal Mummies in Ancient Egypt Cairo: The American University in Cairo Press; 2005. p. 106–19.

4. Zink AR, Sola C, Reischl U, Grabner W, Rastogi N, Wolf H, et al. Characterization of Mycobacterium tuberculosis complex DNAs from Egyptian mummies by spoligotyping. J Clin Microbiol. 2003;41(1):359–67.

5. Zink AR, Grabner W, Reischl U, Wolf H, Nerlich AG. Molecular study on human tuberculosis in three geographically distinct and time delineated populations from ancient Egypt. Epidemiology and Infection. 2003;130(2):239–49.

6. Zink A, Nerlich AG. Molecular analyses of the “Pharaos:” feasibility of molecular studies in ancient Egyptian material. American journal of physical anthropology. 2003;121(2):109–11.

7. Ikram S. Divine Creatures: Animal Mummies in Ancient Egypt. Cairo: Amercain University in Cairo Press; 2005.

8. Hancock JA, Kushlan JA, Kahl MP. Storks, Ibises and Spoonbills of the World. London: Academic Press; 1992.

9. Ray JD. Observations on the Archive of LJor. Journal of Egyptian Archaeology. 1978;64:113–20.

10. Wade AD, Ikram S, Conlogue G, Beckett R, Nelson AJ, Colten R. Backroom Treasures: CT Scanning of Two Ibis Mummies from the Peabody Museum Collection. 2011.

11. Massiera M, Mathieu B, Rouffet F. Speculations on the Role of Animal Cults in the Economy of Ancient Egypt. Apprivoiser le sauvage/Taming the Wild. 2015;11:211–28.

12. Ikram S. An Eternal Aviary, Bird Mummies from Ancient Egypt. In: Bailleul Le-Suer R, editor. Between Heaven and Earth: Birds in Ancient Egypt. Chicago: The Oriental Institute of the University of Chicago; 2012. p. 41–8.

13. Wasef S, Wood R, El Merghani S, Ikram S, Curtis C, Holland B, et al. Radiocarbon dating of Sacred Ibis mummies from ancient Egypt. Journal of Archaeological Science: Reports. 2015;4:355-61.

14. Richardin P, Porcier S, Ikram S, Louarn G, Berthet D. Cats, Crocodiles, Cattle, and More: Initial Steps toward Establishing a Chronology of Ancient Egyptian Animal Mummies. Radiocarbon. 2017;59(2):595–607.

15. Davies S, Smith HS. Sacred Animal Temples at Saqqara. In: Quirke S, editor. The Temple in Ancient Egypt: New Discoveries and Recent Research. London 1997. p. 112–31.

16. Martin GT. The Sacred Animal Necropolis at North Saqqâra: The Southern Dependencies of the Main Temple Complex. London; 1981.

17. Zaghloul HO. Fruhdemotische Urkunden aus Hermupolis. Bulletin of the Center of Papyrological Studies. Cairo1985.

18. Kessler D, Nur el-Din A. Tuna al-Gebel. In: Ikram S, editor. Divine Creatures: Animal Mummies in Ancient Egypt. Cairo: Amercain University in Cairo Press; 2005. p. 120–63.

19. Kessler D, Nur el-Din A. Der Tierfriedhof von Tuna el-Gebel. Antike Welt. 1994;3:252-65.

20. Cuvier G. Discourse of the revolutions on the surface of the earth: determination of the birds called ibis by the ancient Egyptians1825.

21. Kessler D, el-Din AeHN. Tuna al-Gebel. divine creatures: animal mummies in ancient egypt. 2005;7948:120.

22. Meeks D. Les couveuses artificielles en Egypte. Travaux du Centre Camille Jullian. 1997:132–4.

23. Moodie RL. Roentgenologic studies of Egyptian and Peruvian mummies. Field Museum Press. 1931.

24. Dabney J, Knapp M, Glocke I, Gansauge MT, Weihmann A, Nickel B, et al. Complete mitochondrial genome sequence of a Middle Pleistocene cave bear reconstructed from ultrashort DNA fragments. Proc Natl Acad Sci U S A. 2013;110(39):15758–63.

25. Meyer M, Kircher M. Illumina sequencing library preparation for highly multiplexed target capture and sequencing. Cold Spring Harbor protocols. 2010;2010(6):pdb.prot5448.

26. Wasef S, Huynen L, Millar CD, Subramanian S, Ikram S, Holland B, et al., editors. ’Fishing’ for mitochondrial DNA in mummified Sacred Ibis: Development of a targeted enrichment protocol resolves the ancient Egyptian DNA survival debate. the International Symposium of Animals in Ancient Egypt, ISAAE 1; 2016 2018; Lyon, France: Leiden: Sidestone Press.

27. Wasef S, Huynen L, Millar CD, Subramanian S, Ikram S, Holland B, et al. Fishing for Mitochondrial DNA in The Egyptian Sacred Ibis Mummies. bioRxiv. 2018.

28. Li H, Durbin R. Fast and accurate short read alignment with Burrows-Wheeler transform. Bioinformatics. 2009;25(14):1754–60.

29. Li H, Handsaker B, Wysoker A, Fennell T, Ruan J, Homer N, et al. The Sequence Alignment/Map format and SAMtools. Bioinformatics. 2009;25(16):2078–9.

30. Okonechnikov K, Conesa A, Garcia-Alcalde F. Qualimap 2: advanced multi-sample quality control for high-throughput sequencing data. Bioinformatics. 2015.

31. Jonsson H, Ginolhac A, Schubert M, Johnson PL, Orlando L. mapDamage2.0: fast approximate Bayesian estimates of ancient DNA damage parameters. Bioinformatics. 2013;29(13):1682–4.

32. Miller MA, Pfeiffer W, Schwartz T, editors. Creating the CIPRES Science Gateway for inference of large phylogenetic trees. Gateway Computing Environments Workshop (GCE), 2010; 2010: Ieee.

33. Librado P, Rozas J. DnaSP v5: a software for comprehensive analysis of DNA polymorphism data. Bioinformatics. 2009;25(11):1451–2.

34. Nei M. Molecular Evolutionary Genetics. NY: Columbia University Press; 1987.

35. http://www.fluxus-engineering.com/. Network 4.6.1.1. ed: Fluxus Technology Ltd; 2015.

36. Bouckaert R, Heled J, Kuhnert D, Vaughan T, Wu CH, Xie D, et al. BEAST 2: a software platform for Bayesian evolutionary analysis. PLoS Comput Biol. 2014;10(4):e1003537.

37. Bouckaert RR, Drummond AJ. bModelTest: Bayesian phylogenetic site model averaging and model comparison. BMC Evol Biol. 2017;17(1):42.

38. Tamura K, Stecher G, Peterson D, Filipski A, Kumar S. MEGA6: Molecular Evolutionary Genetics Analysis version 6.0. Mol Biol Evol. 2013;30(12):2725–9.

39. Yang Z. PAML 4: phylogenetic analysis by maximum likelihood. Mol Biol Evol. 2007;24(8):1586–91.

40. Kimura M. A simple method for estimating evolutionary rates of base substitutions through comparative studies of nucleotide sequences. Journal of molecular evolution. 1980;16(2):111–20.

41. Paradis E. Analysis of Phylogenetics and Evolution with R. Springer. New York. 2012.

42. Wickham H. ggplot2: an Implementation of the Grammar of Graphics, Version 0.8. 8. 2010.

43. Bandelt H-J, Forster P, Röhl A. Median-joining networks for inferring intraspecific phylogenies. Molecular biology and evolution. 1999;16(1):37–48.

44. Sorenson MD, Quinn TW. Numts: A challenge for avian systematics and population biology. Auk. 1998;115(1):214–21.

45. Carpenter ML, Buenrostro JD, Valdiosera C, Schroeder H, Allentoft ME, Sikora M, et al. Pulling out the 1%: whole-genome capture for the targeted enrichment of ancient DNA sequencing libraries. Am J Hum Genet. 2013;93(5):852–64.

46. Enk JM, Devault AM, Kuch M, Murgha YE, Rouillard J-M, Poinar HN. Ancient Whole Genome Enrichment Using Baits Built from Modern DNA. Molecular Biology and Evolution. 2014;31(5):1292–4.

