## Supplementary data for "Mitogenomic Diversity in Sacred Ibis Mummies sheds light on early Egyptian practices"

### The Sacred Ibis Catacombs

The Egyptian Ministry of State for Antiquity approved sampling of the Sacred Ibis mummies from the main Ibis catacombs (Fig. S1) for the purpose of this research. The permission allowed sampling of Sacred Ibis mummies to be conducted under ‘authorities supervision’ and for dedicated research purposes only. The samples were collected from only three of the main Ibis cemeteries: Saqqara, Tuna El-Gebel, and Abydos, (Fig. S1).

**Fig. S1. The location of the main archaeological sites of ancient Egypt**. The Ibis symbol indicates the location of the main Sacred Ibis catacombs.

### Necropolis at North Saqqara

Saqqara is a village on the west bank of the Egyptian Nile, about 30 km south from the capital city Cairo and is best known for being the site of the Step Pyramid (c. 2550 BC). North of this is the site of one of the largest and the most diverse animal necropolis in Egypt, the Sacred Animal Necropolis (SAN) which is located at the centre of North Saqqara.

The SAN is comprised of a Main Temple complex, with two Ibis catacombs as well as other catacombs for mummified cows, falcons, and baboons (Fig. S2). The discovery of the Sacred Ibis tombs by scholars dates to the 18th century [1]. The catacombs consist of a number of main arterial passageways lined with ‘galleries’, which are rooms on both sides of the passageway. The numerous galleries functioned as storage areas for mummy pots. These were stacked carefully, with a layer of sand separating each layer of vessels. Furthermore, some of the walls in the galleries had a series of ‘niches’ cut to store small limestone chests that housed a mummified bird. This type of burial may be indicative of a special offering from a high-ranking person or a sacred bird burial [1]. A curious feature associated with the temple is a small courtyard garden, planted with bushes, which might have been where the bird that held cult significance was kept. The priests of Thoth[2] were to look after these temple birds [3].

**Fig. S2.** **Details of the SAN (Sacred Animal Necropolis) at Saqqara**. Original drawing by J. Hodges edited by Miriam Alexander.

### Tuna al-Gebel

Tuna al-Gebel, also known as Hermopolis Magna by the ancient Greeks, is located on the west side of the Nile, close to the desert margin. The southern part of the site contains a vast animal necropolis containing several million mummified Sacred Ibis, a significant number of baboons, as well as some other mummified animals Both ibis and baboons represented the god Thoth and were buried together in subterranean galleries starting in c. 664 BC and continuing through the Ptolemaic era into Roman times (possibly until c. AD 200).

The catacombs are estimated to contain more than 4 million Sacred Ibis mummies. Texts show that an ‘Ibiotropheion’, the Greek name for a special place for breeding ibis, existed at Tuna al-Gebel [4]. These represent an ideal opportunity to study both the evolution of animal cemeteries and related ritual beliefs over more than 700 years of ancient Egyptian history. The gallery (Fig. S3) from which our sample was taken was constructed during the 26th Dynasty (c. 685–525 BC) and had been cut in a westward direction. A long passage stretching from north to south is filled with pottery jars that contain Sacred Ibis mummy bundles. It is entered from the north. The main subterranean passage had rock cut chambers along its sides. The jars of that period were wide-mouthed and closed with plastered linen [4].

**Fig. S3. Layout of the Sacred Ibis catacomb (Ibiotropheion) at Tuna el-Gebel**. Showing the different galleries. The original map was supplied by courtesy of Dieter Kessle.

### Abydos

Abydos is one of the most ancient cities of Upper Egypt and is the burial place for its earliest rulers. It is about 11 kilometres west of the river Nile at latitude 26°10' N, near the modern towns of el-'Araba el Madfuna and al-Balyana, and south of the city of Sohag. The city was known as Abdju in hieroglyphics (3bdw or AbDw), which means "the hill of the symbol or shrine". There are several ibis burial places scattered all over the site of Abydos [5] Ibis cemetery, which is located near Umm el-Qa′ab(Fig. S4) containing not only mummified ibis, but also human burials [6].Other denser Ibis burials in the Abydos region are found in Shunet el-Zebib; the funerary enclosure for King Khasekhemwy (2690 BC). These burials were used as Ibis cemeteries from the 22nd Dynasty (945 BC – 715 BC) onward [7].

**Fig. S4. Details of the cemetery at Abydos**. Showing the location of the Ibis cemetery.

**References**

1.	Nicholson PT. The Sacred Animal Necropolis at North Saqqara. Divine creatures: Animal mummies in ancient Egypt. 2005;7948:44.

2.	Martin GT. The Sacred Animal Necropolis at North Saqqâra: The Southern Dependencies of the Main Temple Complex. London; 1981.

3.	Ray JD. Observations on the Archive of Ḥor. Journal of Egyptian Archaeology. 1978;64:113–20.

4.	Kessler D, Nur el-Din A. Tuna al-Gebel. In: Ikram S, editor. Divine Creatures: Animal Mummies in Ancient Egypt. Cairo: Amercain University in Cairo Press; 2005. p. 120-63.

5.	Ikram S. Animals in the Ritual Landscape at Abydos. American University in Cairo; 2007.

6.	Peet TE. The Cemeteries of Abydos: Part 2. 1911-1912: Adegi Graphics LLC; 1999.

7.	Ayrton ER, Currelly CT, Weigall AE. Abydos III. EEF, London. 1904.

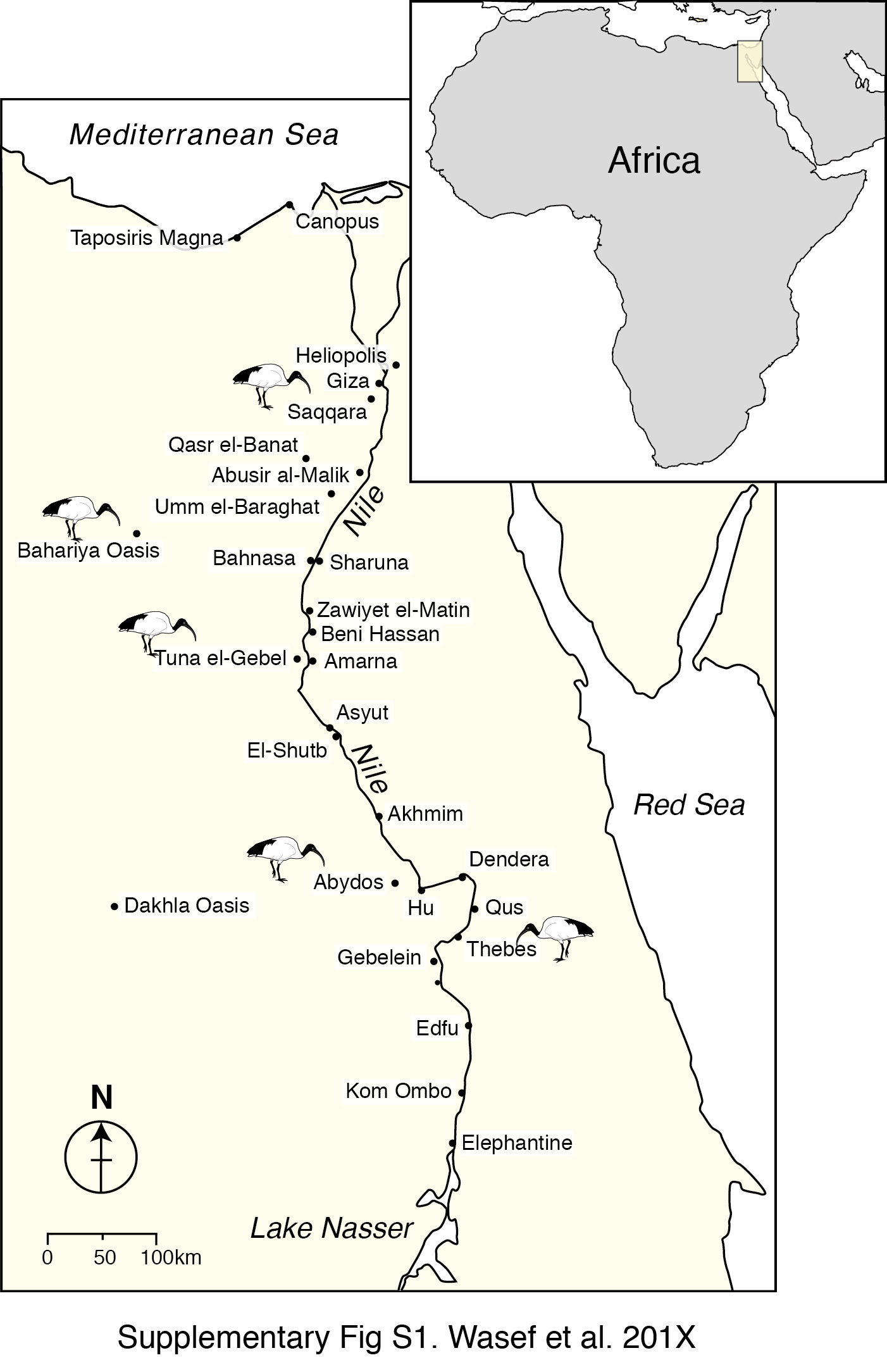

Fig. S1.

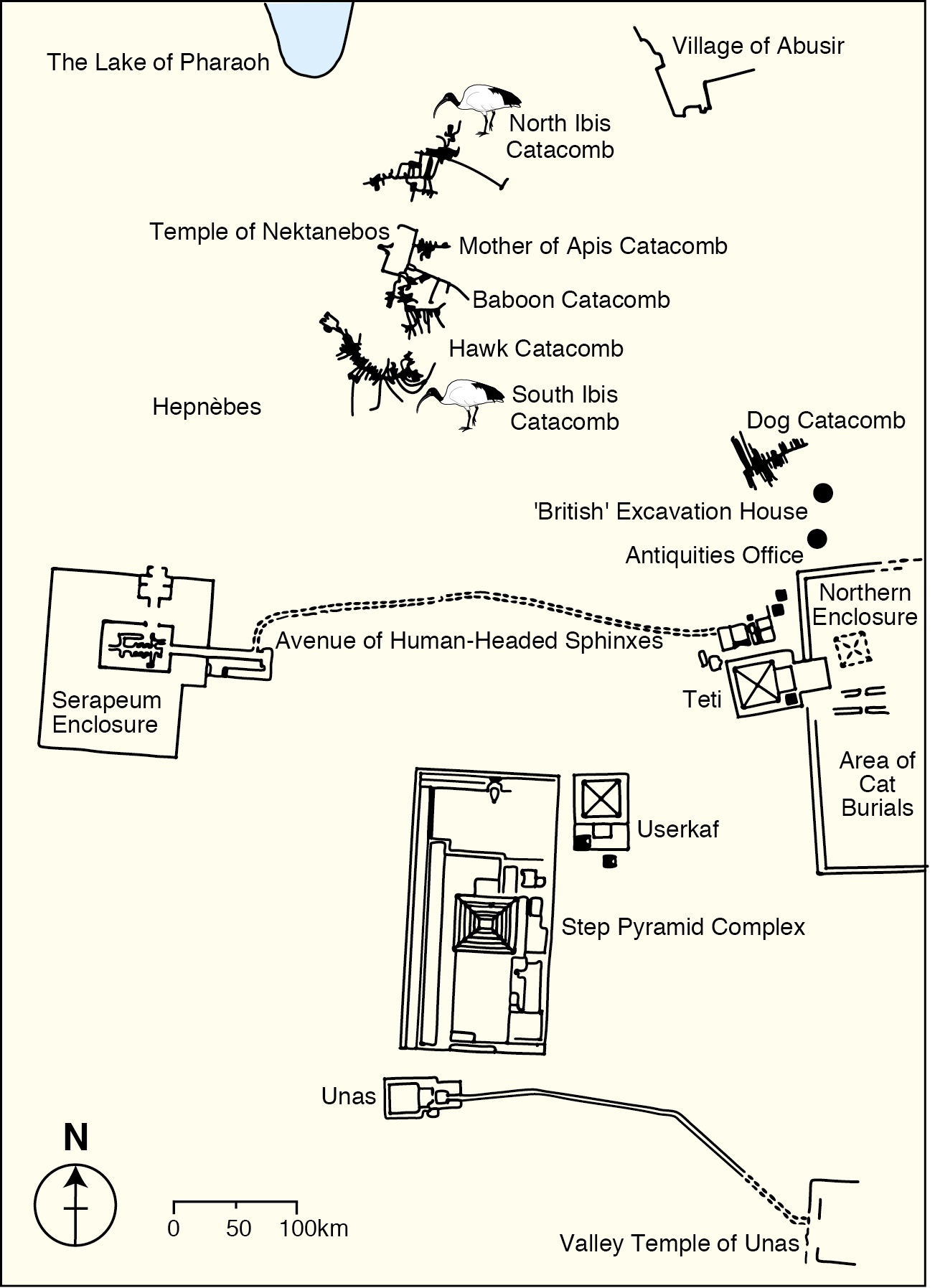

Fig. S2.

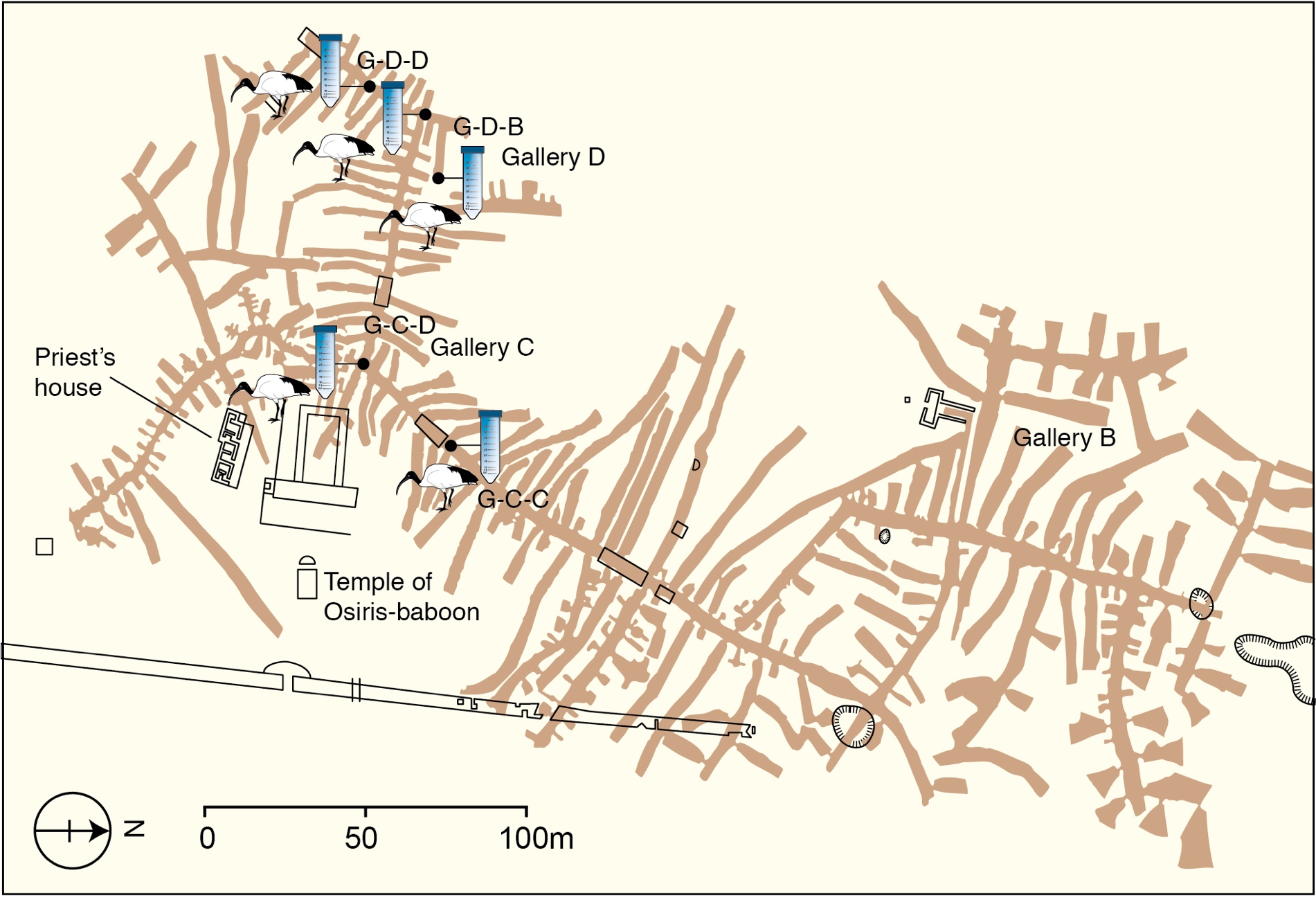

Fig. S3.

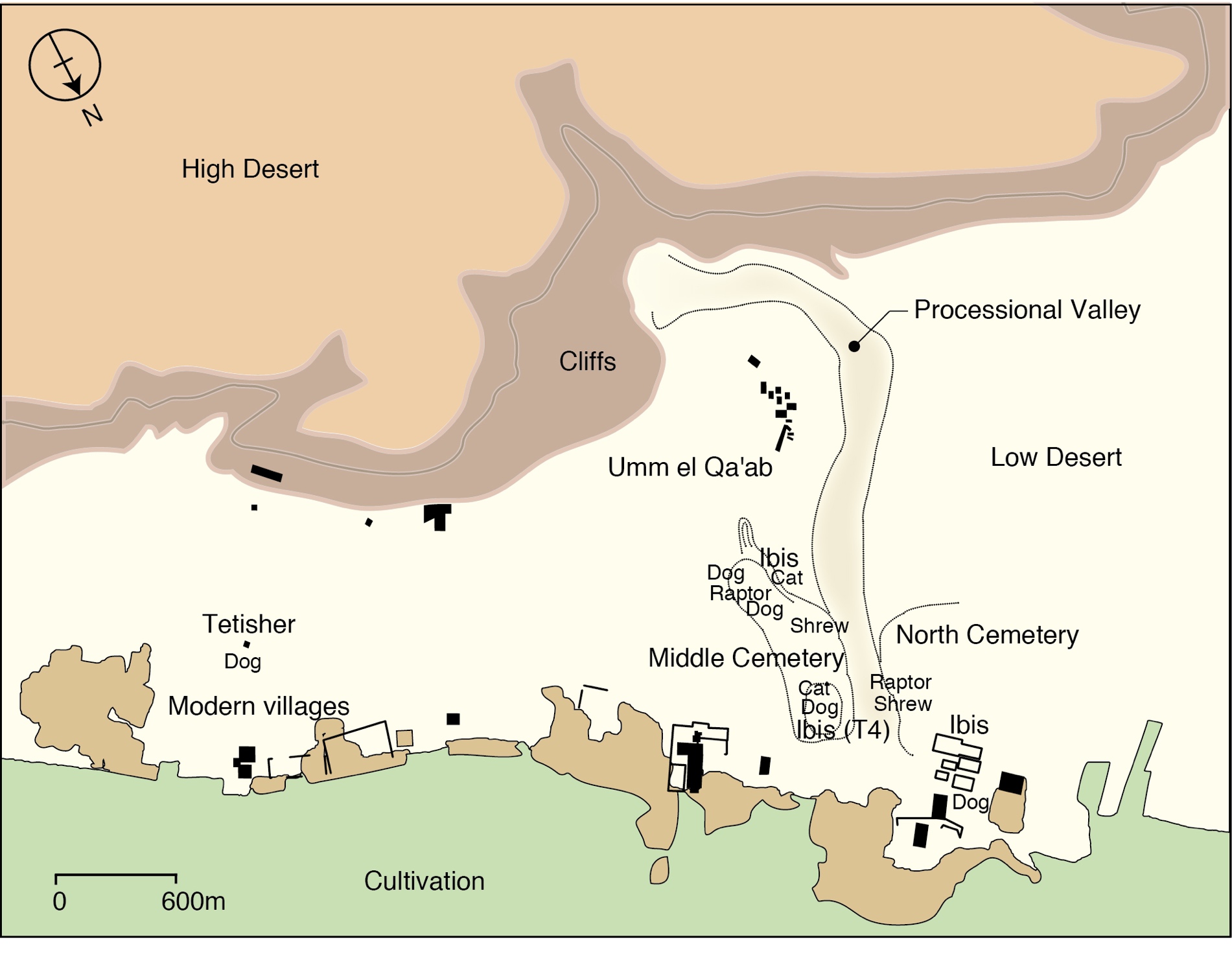

Fig. S4.

Table S1 | Description of data generated using the Modern and Ancient Sacred Ibis samples showing; Shotgun sequencing results for unique mitochondrial endogenous content, coverage and clonality; The target capture hybridisation results for unique mitochondrial sequences, coverage, enrichment and clonality. ‘Total reads’ refers to the sequence data generated before aligning to the Sacred Ibis mitochondrial genome. ‘Unique’ refers to the fraction of mapped mitochondrial reads after removing the clonal sequences. The enrichment rate (x fold) is calculated by dividing the % of unique endogenous mitochondrial sequences after capture by the total number of shotgun sequences. Since, the nuclear reference modern Ibis genome is not available, we could only calculate the levels using the known Sacred Ibis mitogenome. Hence the nuclear reads in the raw data could not be included in calculating the endogenous DNA levels.

|  | **Direct sequencing** | | | | **Target capture** | | | | | **Unique enrichment (x)** |
| --- | --- | --- | --- | --- | --- | --- | --- | --- | --- | --- |
| **Sample** | **Total reads** | **Unique mapped** | **Mean read length** | **mean coverage** | **Total reads** | **Unique mapped** | **Mean read length** | **Mean coverage(x)** | **Temperature (C°)** |  |
| **Tuna_1** | 47,428,937 | 22 | 60.1 | 0.06 | 60,239,019 | 701 | 73.8 | 1.94 | 45 | 25 |
| **Thebes_1** | 30,552,363 | 68 | 44.4 | 0.16 | 32,645,320 | 1,301 | 46.6 | 3.29 | 55 | 18 |
| **Thebes_1** | 27,975,319 | 13 | 49.9 | 0.04 | 27,677,743 | 346 | 53.4 | 0.9 | 55 | 27 |
| **Thebes_2** | 17,681,629 | 32 | 45.9 | 0.07 | 16,595,500 | 212 | 53.3 | 0.53 | 55 | 7 |
| **Tuna_2** | 20,711,640 | 377 | 45.6 | 0.86 | 20,817,703 | 5,668 | 47.7 | 14.05 | 55 | 15 |
| **Thebes_2** | 7,539,230 | 17 | 45.9 | 0.07 | 3,857,563 | 1,108 | 77.6 | 3.34 | 57 | 127.4 |
| **Thebes_3** | 20,665,829 | 793 | 66.6 | 2.35 | 839,683 | 11,401 | 73.3 | 37.62 | 57 | 354 |
| **KomOmbo** | 6,193,092 | 238 | 77.8 | 0.63 | 914,909 | 8,781 | 75.9 | 22.86 | 57 | 250 |
| **Roda** | 2,322,995 | 36 | 90 | 0.09 | 5,776,199 | 4,810 | 69.3 | 13.83 | 57 | 54 |
| **Sohag_2** | 63,609,333 | 1,177 | 53.9 | 4.13 | 5,222,068 | 23,361 | 92.3 | 121.88 | 57 | 242 |
| **Sohag_3** | 61,488,812 | 144 | 67.7 | 0.57 | 1,911,606 | 21,063 | 87 | 82.63 | 57 | 4705 |
| **Sohag_1** | 22,790,097 | 2,236 | 44.9 | 5.61 | 13,321,380 | 962 | 43 | 2.73 | 65 | 1 |
| **Sohag_1** | 6,749,628 | 2,870 | 45.6 | 6.84 | 26,119,217 | 3,249 | 46.2 | 10.4 | 65 | 0.29 |
| **Saqara_14** |  |  |  |  | 3,551,057 | 1,048 | 84.7 | 4.27 | 57 |  |
| **Saqara_15** |  |  |  |  | 2,493,552 | 628 | 77.6 | 3 | 57 |  |
| **Saqara_16** |  |  |  |  | 1,458,625 | 509 | 92.1 | 2.26 | 57 |  |
| **Saqara_33** |  |  |  |  | 1,882,473 | 990 | 94.8 | 5 | 57 |  |
| **South Africa Population 1 _1** | 11,786,439 | 98 | 59.9 | 0.51 | 18,202,562 | 9,510 | 89.9 | 55.15 | 65 | 63 |
| **South Africa Population 1_2** | 33,360,459 | 59 | 53.1 | 0.29 | 35,400,567 | 19,936 | 89.9 | 116.81 | 65 | 318 |
| **South Africa Population 1 _3** | 206,187,225 | 6,663 | 93.2 | 36.43 | 21,488,008 | 15,899 | 91.8 | 92.75 | 65 | 23 |
| **South Africa Population 2_1** | 46,354,443 | 58 | 50.6 | 0.26 | 22,291,370 | 15,129 | 88.7 | 88.05 | 65 | 542 |
| **South Africa Population 2_2** | 7,061,662 | 28 | 73.3 | 0.11 | 21,232,888 | 4,207 | 89.1 | 23.49 | 65 | 50 |
| **Kenya_F1** | 5,613,148 | 1,900 | 110.9 | 12.59 | 6,485,517 | 31,331 | 116.5 | 259.22 | 60 | 14 |
| **Kenya_F3** | 751,779 | 1,034 | 92.1 | 6.1 | 3,536,543 | 26,489 | 100.5 | 206.06 | 60 | 5 |
| **Kenya_F4** | 2,280,496 | 677 | 103.6 | 3.86 | 4,491,818 | 25,102 | 108.5 | 193.3 | 60 | 19 |
| **Kenya_F5** | 1,586,081 | 25 | 100.3 | 0.1 | 2,380,562 | 3,673 | 105.9 | 24.42 | 60 | 98 |
| **Kenya_F6** | 1,565,081 | 780 | 102 | 4.1 | 3,634,734 | 28,280 | 112.7 | 232.89 | 60 | 16 |
| **Kenya_F7** | 1559235 | 956 | 112.1 | 6.62 | 2,658,086 | 29,507 | 117.3 | 243.32 | 60 | 18 |
